# Sex-biased Fibroblast Subpopulations and Transcriptional Programs Reveal Mechanisms of Skin Lesion Development in Systemic Sclerosis

**DOI:** 10.64898/2026.05.06.723148

**Authors:** Chiranan Khantham, Inmaculada Rodriguez-Martin, Martin Kerick, Gonzalo Villanueva-Martin, José Luis Callejas, Norberto Ortego-Centeno, Alfredo Guillen-Del-Castillo, Carmen P. Simeón-Aznar, Ricardo Ruiz-Villaverde, Eduardo Andrés-León, Javier Martin, Marialbert Acosta-Herrera

## Abstract

Systemic sclerosis (SSc) is an autoimmune connective tissue disease with pronounced sex differences: females are more frequently affected and males develop more severe skin fibrosis. The cellular mechanisms of this disparity remain unclear. Here we use single-cell transcriptomics of lesional, non-lesional, and healthy skin to define fibroblast states and sex-biased transcriptional programs during lesion development. We identify a sex-dependent divergence in SSc fibrotic regulation. Female fibroblasts exhibit heightened inflammatory signaling and canonical TGF-β-driven extracellular matrix production, whereas male fibroblasts preferentially engage non-canonical TGF-β pathways, mechanotransduction, and MYC-associated stress programs. We further reveal that the fibrotic lesional environment shows sex differences: *SFRP2⁺DPP4⁺* fibroblasts predominate in females and *COL11A1⁺/COCH*⁺ in males. Our findings uncover cellular mechanisms underlying sex differences in SSc fibrosis, highlight opportunities for sex-informed therapeutic strategies and underscore the necessity of integrating biological sex into precision medicine frameworks to identify divergent molecular drivers of fibrotic disease.

## Introduction

Systemic sclerosis (SSc) is a complex autoimmune disease characterized by autoimmunity, progressive fibrosis of the skin and internal organs, and vasculopathy^1^. Skin involvement is a defining clinical feature of SSc and serves as the primary criterion for classifying the disease into limited cutaneous (lcSSc) and diffuse cutaneous (dcSSc) subtypes^2^. Despite its low prevalence, SSc carries one of the highest mortality rates among rheumatic diseases and imposes substantial clinical and socioeconomic burdens^1^.

A hallmark of SSc, as in many autoimmune diseases, is its pronounced sex disparity. Females are predominantly affected, with a female-to-male ratio of 8:1^3^, whereas males more frequently develop dcSSc and exhibit more severe organ involvement and poorer prognosis^4^. These differences have been attributed to the interplay of sex chromosome-linked effects, hormonal influences, and immune function^3–5^. However, these mechanisms do not fully capture the complexity of sex differences in SSc. Genetic susceptibility has been extensively studied through genome-wide association studies (GWAS), which have identified multiple susceptibility loci implicating immune and fibrotic pathways^6^. Recent sex-stratified GWAS have revealed sex-specific genetic associations, uncovering previously unrecognized regulatory mechanisms that may drive sex-biased transcriptional programs^7^. Together, these findings position sex as a key determinant of SSc susceptibility, clinical manifestation, and disease progression. Despite these advances, the mechanisms underlying sex bias in SSc remain poorly defined and warrant further investigation.

Fibroblasts are central mediators of fibrosis, producing excessive collagen and extracellular matrix (ECM) components that result in thickened, stiff, and hardened skin. In SSc, persistent fibroblast-to-myofibroblast transition, driven primarily by TGF-β signaling and mechanical stress^8^, promotes the accumulation of contractile myofibroblasts, whose abundance correlates with skin fibrosis severity and modified Rodnan skin score (MRSS), a clinical measure of skin fibrosis^9^. Fibroinflammatory skin involvement remains difficult to treat, and current therapies cannot reverse established fibrosis^2^. Although agents targeting fibroblasts or immune–fibroblast pathways are in clinical use or clinical trials^2,10^, their efficacy remains limited without a detailed understanding of the sex-differentiated mechanisms.

Single-cell RNA sequencing (scRNA-seq) has revealed substantial fibroblast heterogeneity in SSc skin^11–14^. However, sex-biased effects in scleroderma fibroblasts remain largely unexplored. To address this gap, we generated a comprehensive single-cell transcriptomic atlas of lesional, non-lesional, and healthy skin from SSc patients and controls, focusing on fibroblast lineages to define sex-biased transcriptional programs, fibroblasts states and sex-differentiated pathways. Our analysis uncovers distinct mechanisms of fibrogenesis, with inflammatory and growth factor-driven programs predominating in females, while males exhibit enhanced disease severity associated with non-canonical TGF-β signaling, mechanotransduction, and MYC-driven metabolic and stress responses, providing a cellular basis for sex differences in SSc and supporting the development of sex-informed therapeutic strategies.

## Results

### Single-cell profiling of SSc and healthy skin

A total of 14 transcriptionally distinct cell populations were identified (Fig. 1a), capturing the major structural and immune compartments of the skin. Prominent populations included fibroblasts, vascular endothelial cells, and T cells, along with additional specialized tissue-resident populations. These observed cell populations were largely consistent with those reported in previous single-cell studies of human skin^11,12,15^. The composition of these cell clusters was consistent across sex, disease status, and different skin types (Fig. 1b–d). A dot plot showing canonical marker genes used to define each major skin cell type is shown in Fig. 1e.

**Fig. 1.**
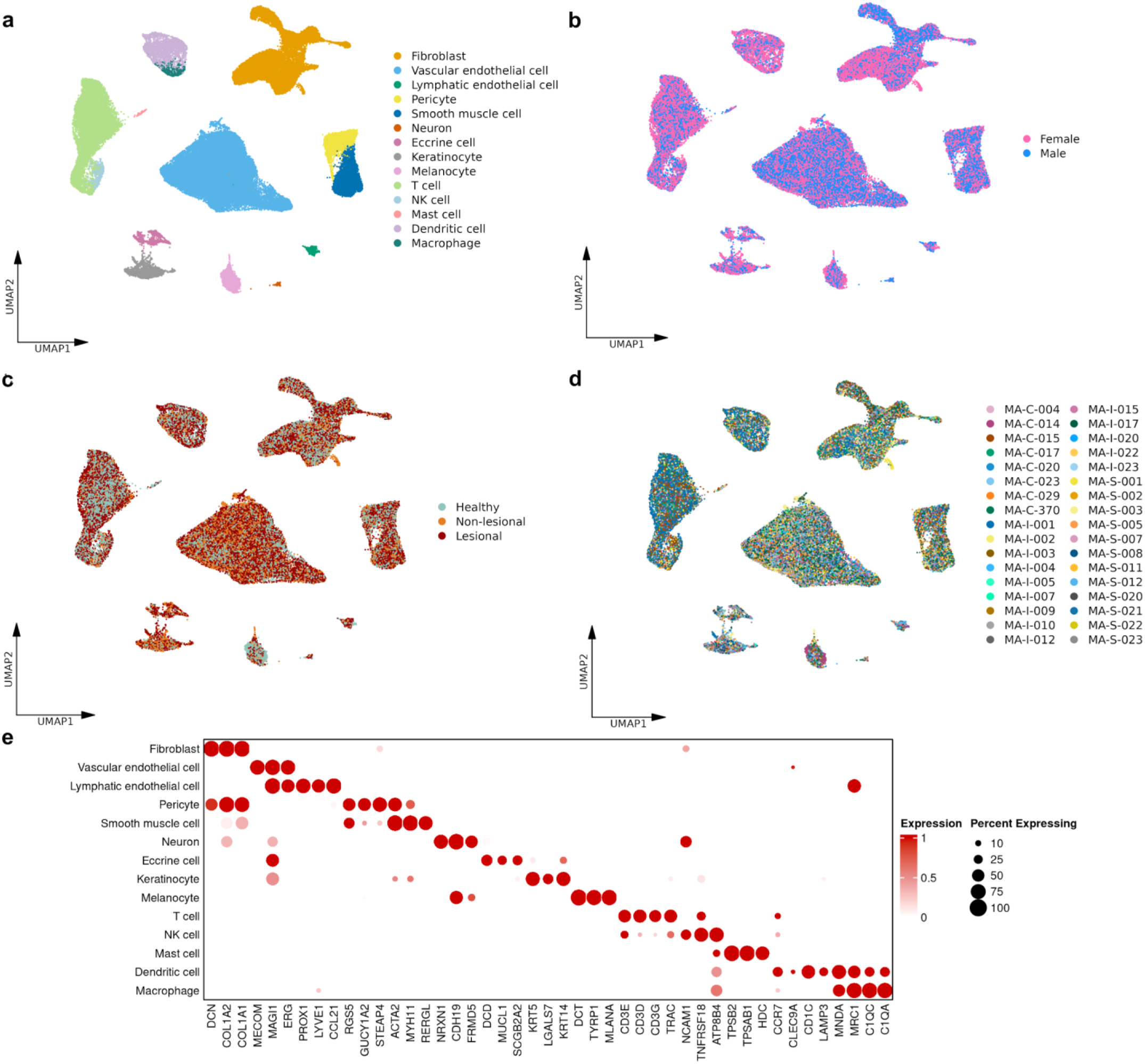
Single-cell landscape of SSc and healthy skin. **a** UMAP visualization of 14 skin-derived cell types profiled by scRNA-seq, coloured by annotated cell types. **b** UMAP showing cells coloured by sex. **c** UMAP showing cells coloured by clinical skin status. **d** UMAP showing cells coloured by sample identity. **e** Dot plot depicting the expression of canonical marker genes used for cell-type annotation. Colour intensity represents the scaled average expression level, and dot size indicates the percentage of cells within each cluster expressing the specified marker.

### Sex-biased transcriptional programs in fibroblasts drive divergent SSc lesion development

Fibroblasts are the key effector cells driving fibrotic remodeling in SSc, and their persistent activation leads to the expansion of myofibroblasts^8^. To elucidate the sex-biased transcriptional programs governing SSc lesion formation, we applied a *sex × skin types* interaction model (contrast defined in Methods) to pseudobulk fibroblast gene expression profiles, identifying sex-modulated gene sets and pathways in fibrogenesis (Supplementary Data 1). These findings were complemented by sex-stratified lesional versus non-lesional analyses to resolve the directionality of these activation states (Supplementary Data 2-3; Supplementary Fig. 1).

In total, we identified 533 sex-biased differentially expressed genes (DEGs) through the interaction contrast (false discovery rate [FDR] < 0.05; absolute log_2_-fold change [|logFC|] > 0.5). Interestingly, 135 genes were significantly altered during lesion development in females and 398 exhibited significant transcriptional changes in males (Fig. 2a). *EPAS1*, a hypoxia-inducible factor, was the most significant female-biased DEG (logFC = 1.41). Conversely, *S100A4*, a critical damage-associated molecular pattern in SSc fibrosis^16^, emerged as the top male-biased DEG (logFC = −1.25).

**Fig. 2.**
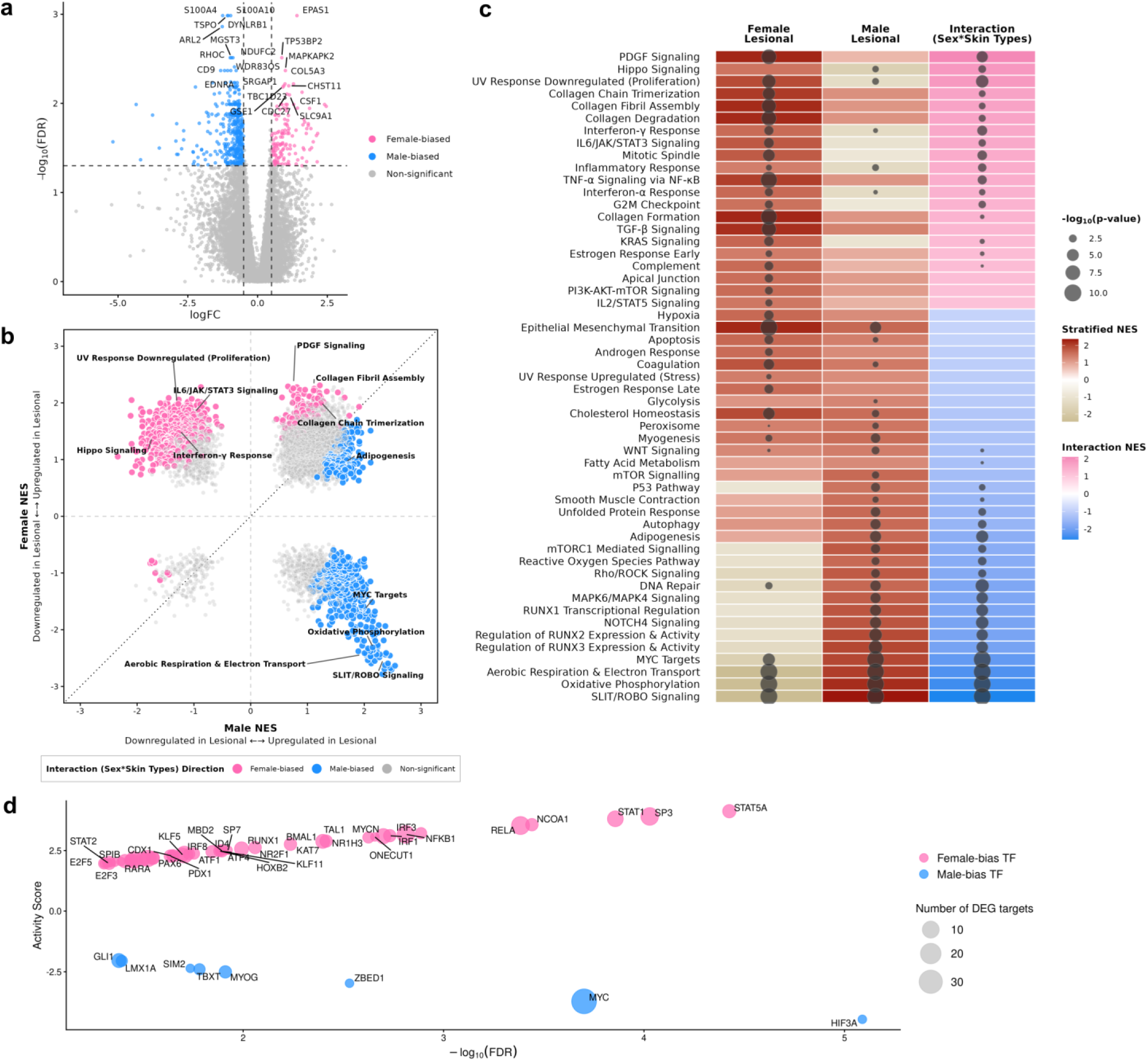
Sex-biased transcriptional programs and transcriptional regulation in fibroblasts during lesional development. **a** Volcano plot showing sex-biased differentially expressed genes (DEGs) in fibroblasts during lesional development, derived from the *sex × skin types* interaction in pseudobulk analysis. Sex-biased DEGs were identified using thresholds of false discovery rate (FDR) < 0.05 and absolute log_2_-fold change (|logFC|) > 0.5, where positive logFC value indicates female-biased transcriptional shifts and negative logFC value indicates male-biased transcriptional shifts. **b** Scatter plot comparing pathway enrichment scores (normalized enrichment score, NES) between females (y-axis) and males (x-axis) from stratified pseudobulk analyses comparing lesional versus non-lesional skin. Pathways are coloured by sex bias (female-biased, pink; male-biased, blue; non-significant, grey). Selected pathways are annotated. **c** Dot plot heatmap summarizing pathway enrichment across sex-stratified (lesional versus non-lesional skin) and interaction (*sex × skin types*) pseudobulk analyses. Colour intensity represents stratified NES (upregulated in lesional skin: red; downregulated: brown) or interaction NES (female-biased: pink; male-biased: blue). Dot size indicates statistical significance; absence of a dot indicates non-significance. **d** Inferred transcription factor (TF) activity derived from sex-biased DEGs using the CollecTRI regulatory network. Positive TF activity scores indicate female-biased TF activity, whereas negative scores indicate male-biased TF activity. Dot size corresponds to the number of differentially expressed target genes regulated by each TF.

Gene set enrichment analysis (GSEA) confirmed that this sex-transcriptional asymmetry reflects a divergence in the pathological programs driving SSc fibrosis (Fig. 2b-c; Supplementary Data 4). While both sexes converged on core fibrotic processes, including epithelial-to-mesenchymal transition and coagulation, the underlying regulatory programs were highly sex-biased, with several pathways exhibiting opposite directionality. Female-biased activation was significantly enriched for Hippo signaling and profibrotic growth factor signaling, specifically the platelet-derived growth factor (PDGF) pathway and its downstream Kirsten rat sarcoma virus (KRAS) signaling node. This was accompanied by a broad suite of immune–inflammatory pathways, including Toll-like receptor (TLR), interleukin-6/JAK/STAT3, interferon, and tumor necrosis factor (TNF), alongside programs governing cycle progression, estrogen response and ECM organization.

In males, we identified several antagonistically regulated pathways, including SLIT/ROBO signaling, MYC targets, and the tumor protein p53 pathway, which have not been previously described in scleroderma fibrogenesis. Males were also specifically enriched for mechanotransduction programs, including smooth muscle contraction and Rho-associated protein kinase (Rho/ROCK) signaling, as well as pathways involving protein homeostasis, DNA repair, mechanistic target of Rapamycin complex 1 (mTORC1), and RUNX-mediated transcriptional programs. Furthermore, a male-specific metabolic shift was evident through the enrichment of adipogenesis, aerobic respiration, electron transport and oxidative phosphorylation, indicating a heightened bioenergetic demand within the male fibrotic niche.

### Sex-biased transcription factor (TF) activity in fibroblasts during SSc lesion development

To identify the regulatory drivers of sex differences in SSc fibrogenesis, we performed TF activity inference. We identified 54 TFs with significant activity changes during lesion development (Fig. 2d; Supplementary Data 5), comprising 42 female-biased and 12 male-biased TFs. The majority of these TFs were not differentially expressed at the mRNA level, indicating that inferred activity was primarily driven by coordinated changes in their target gene expression rather than TF abundance.

Among the female-biased TFs, STAT5A displayed the highest TF activity score, with its regulons enriched for genes involved in inflammatory signaling and epigenetic regulation. Furthermore, STAT1, RUNX1, and multiple interferon regulatory factor (IRF) family members, well-known TFs implicated in systemic autoimmunity^11,17^, were preferentially active in female fibrotic responses. Approximately 45% of female-biased TFs targeted *CREBBP*, an epigenetic regulator essential for myofibroblast differentiation^30^ (Supplementary Data 5). VGLL3 (a master regulator of female-biased inflammatory genes in skin autoimmunity^18^) was not captured by TF activity inference due to its absence from the CollecTRI network database; however, it was significantly upregulated at the mRNA level in female lesional fibroblasts compared with non-lesional fibroblasts (Supplementary Data 2). VGLL3 was predicted to interact with several identified female-biased TFs (Supplementary Fig 2), suggesting that it functions as a female-specific co-factor reinforcing immune–fibrotic regulatory programs in female SSc.

On the other hand, MYC emerged as a central male-biased regulator. Consistent with the pathway enrichment analysis described above, MYC coordinated transcriptional programs associated with fibroblast-to-myofibroblast activation and cellular persistence. Its regulons were enriched for genes involved in cytoskeletal and contractile functions, ECM remodeling, ribosomal biogenesis, RNA processing, cellular metabolism, anti-apoptotic and survival genes (Supplementary Data 5).

### Fibroblast subpopulations in SSc and healthy skin

We identified six transcriptionally distinct fibroblast subtypes across SSc and healthy skin, defined by the following marker signatures: *COCH⁺*, *APOE⁺CXCL12⁺*, *HHIP⁺*, *COL11A1⁺*, *SFRP2⁺DPP4⁺*, and *CLDN1⁺* (Fig 3a-b; Supplementary Data 6). Although these subpopulations were present in both SSc and healthy samples, we observed pronounced shifts in their proportions and transcriptional states associated with the SSc lesional niche, as well as sex-biased distribution patterns. These clusters were consistent with previously reported skin fibroblast populations, recapitulating papillary, reticular, mesenchymal, and pro-inflammatory states^11,14^.

**Fig. 3.**
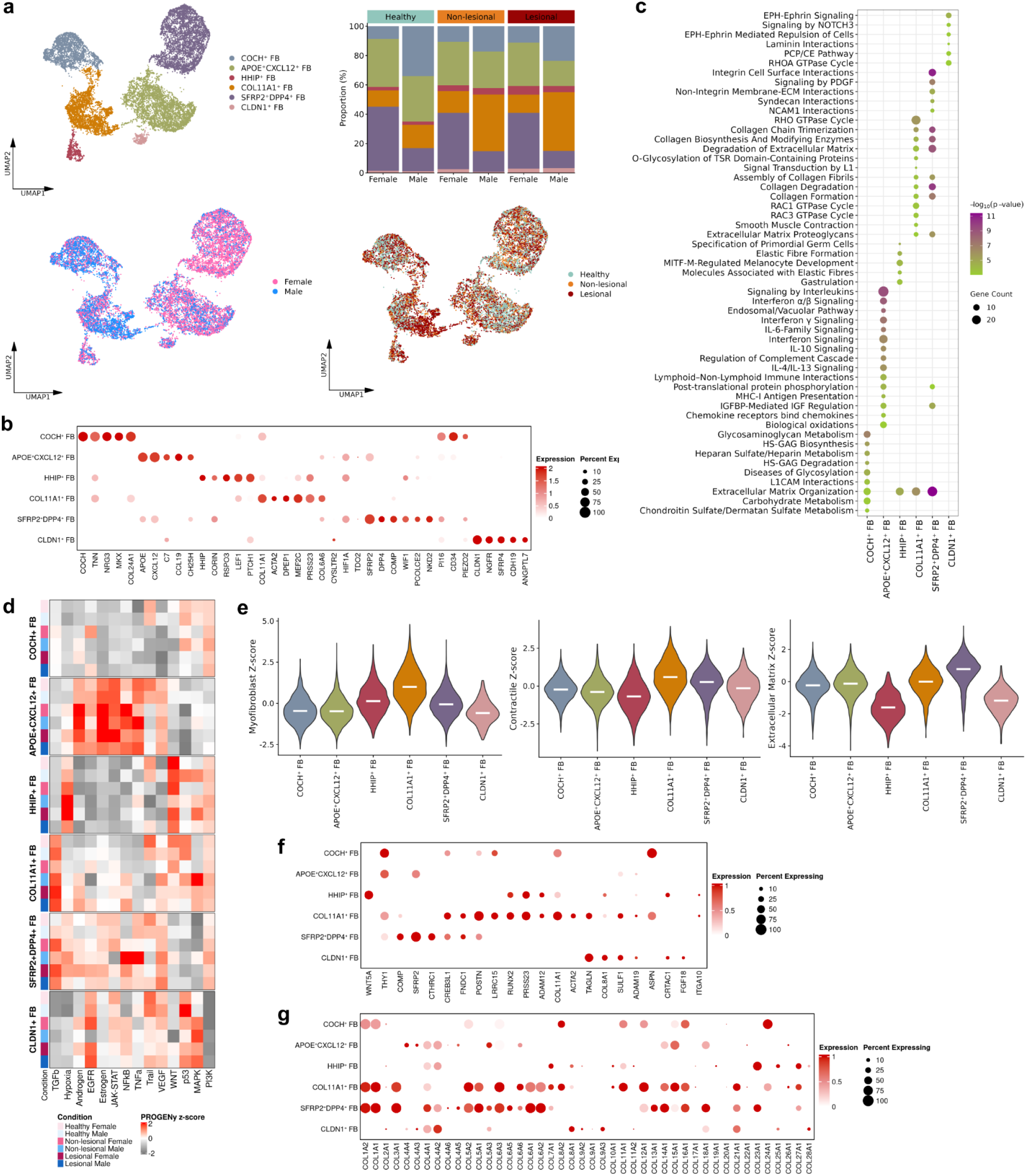
Characterization of fibroblast subpopulations and their functional programs in SSc and healthy skin. **a** UMAP visualization of six fibroblast subtypes, coloured by subtype identity, sex, and skin status, alongside stacked bar plots showing the relative proportions of each fibroblast subtype stratified by sex and skin status. **b** Dot plot of marker genes defining each fibroblast subtype. **c** Bubble plot showing significantly enriched Reactome pathways across fibroblast subtypes identified by over-representation analysis. Colour intensity indicates the level of statistical significance, and circle size represents the number of genes contributing to each pathway. **d** Heatmap of inferred pathway activities across fibroblast subtypes, skin types, and sexes using PROGENy. Colour intensity represents the calculated *Z*-score, where positive values indicate pathway activation and negative values indicate inhibition. **e** Violin plots showing module *Z*-scores for myofibroblast, contractile, and extracellular matrix programs across fibroblast subtypes. **f** Dot plot showing myofibroblast marker gene expression. **g** Dot plot showing collagen-associated gene expression. For all dot plots, colour intensity indicates scaled expression and dot size indicates the percentage of expressing cells.

*COL11A1⁺*and *SFRP2⁺DPP4⁺* subpopulations were identified as the principal drivers of the fibrotic program, both showing marked enrichment for ECM organization and collagen biosynthesis pathways (Fig. 3c). Pathway activity inference confirmed robust activation of TGF-β signaling in both populations across lesional and non-lesional SSc skin (Fig. 3d). *COL11A1⁺* fibroblasts represented a mesenchymal population highly enriched for myofibroblast-associated transcriptional programs (Fig 3e-f). These cells expressed mesenchymal-associated markers (*COL21A1*, *POSTN*)^19^ and myofibroblast markers, most notably *ACTA2*, which encodes α-smooth muscle actin (α-SMA). Functionally, *COL11A1⁺* fibroblasts were uniquely enriched for mechanosensitive signaling pathways, including Rho/Rac GTPase signaling, smooth muscle contraction, and L1-mediated signal transduction^8,20^ (Fig. 3c), and displayed the highest myofibroblast and contractile module scores (Fig 3e).

In contrast, *SFRP2⁺DPP4⁺* cluster represented a transcriptional continuum of papillary and reticular fibroblasts and exhibited the highest ECM score, along with expression of a diverse repertoire of collagen genes (Fig 3e,g). The remaining subpopulations occupied specialized dermal niches. *APOE⁺CXCL12⁺* subpopulation represented a pro-inflammatory state, expressing fibroblast reticular cell–like genes (*CCL19, CH25H*)^14^, and showed enrichment of immune-related pathways (Fig 3c). Finally, we identified hair follicle-associated *COCH⁺* and *HHIP⁺* fibroblasts and a minor *CLDN1⁺* subpopulation displaying Schwann cell–like features.

### Sex-biased fibroblast abundance in SSc lesional skin

To investigate sex-biased compositional shifts in fibroblast subtypes in SSc lesional skin, we performed a neighborhood-based differential abundance analysis using MiloR. In healthy skin, we observed broad and pronounced sex-biased distributions across nearly all fibroblast clusters, with the exception of the *CLDN1⁺* population, establishing a baseline of significant sex differences in dermal fibroblast composition (Fig. 4a).

**Fig. 4.**
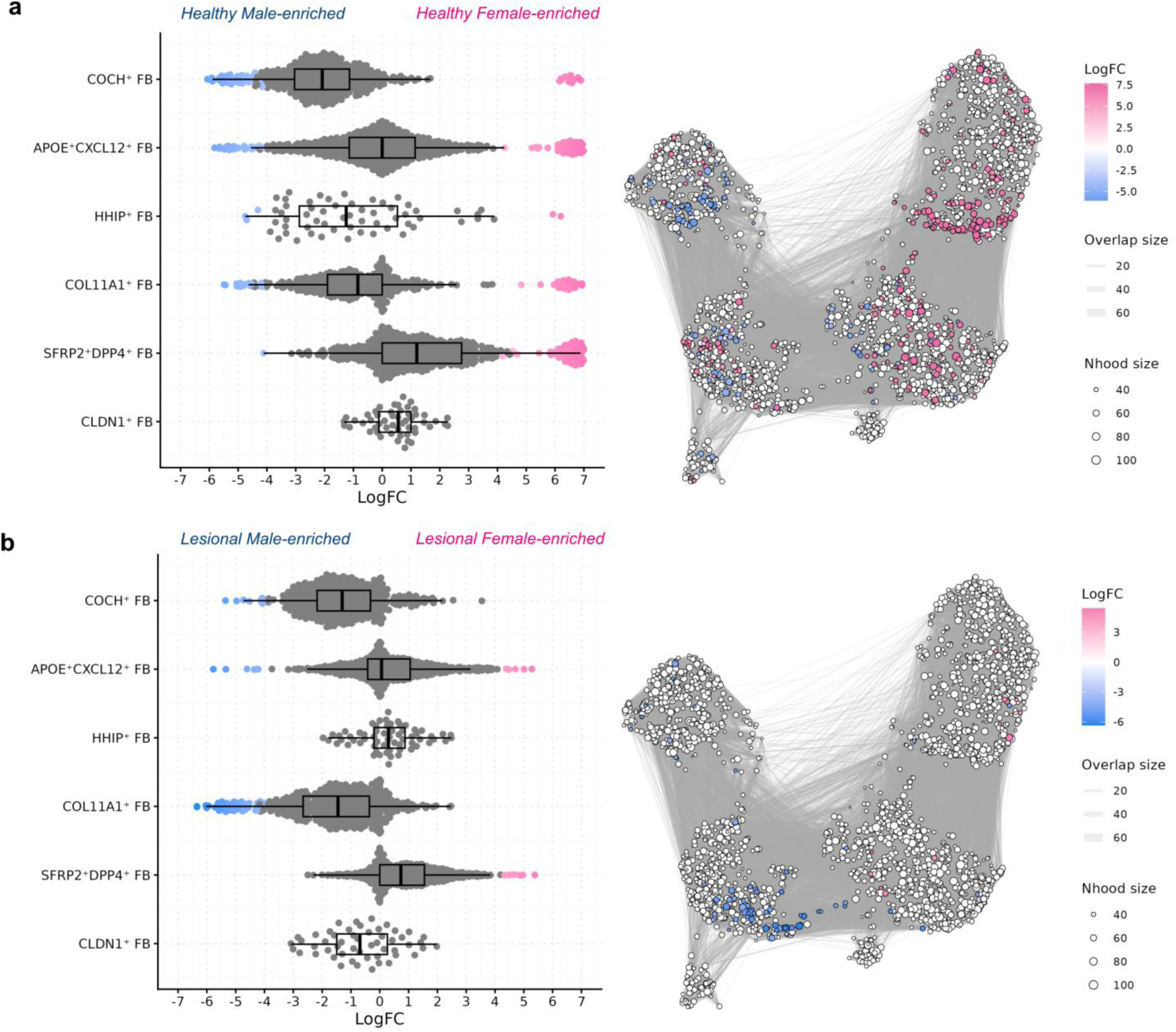
Differential abundance of fibroblast subtypes reveals sex-biased enrichment in lesional and healthy skin. **a-b** Beeswarm plots (left) and neighbourhood UMAPs (right) showing sex-biased differential abundance (DA) of fibroblast subpopulations in healthy (**a**) and lesional (**b**) skin. DA was assessed using Milo; significant neighbourhoods (spatial false discovery rate < 10%) are highlighted. Positive log-fold change (logFC) indicates female enrichment (pink) and negative logFC indicates male enrichment (blue). In beeswarm plots, dots represent neighbourhoods; boxes indicate median and interquartile range. In UMAPs, nodes represent neighbourhoods and edges denote shared cells; node size corresponds to neighbourhood (Nhood) size.

In contrast, sex-differences in SSc lesional skin were limited and largely confined to specific fibroblast subsets associated with fibrotic remodeling (Fig. 4b). These findings indicate that the lesional environment is dominated by strong profibrotic programs, which concentrate sex-biased effects within key pathogenic fibroblast lineages. Specifically, the *SFRP2⁺DPP4⁺* cluster was selectively enriched in female lesional skin, whereas *COL11A1⁺* and *COCH⁺* subpopulations were preferentially expanded in male SSc lesional skin. The pro-inflammatory *APOE⁺CXCL12⁺* subpopulation displayed a mosaic pattern, with distinct neighborhoods enriched in either sex.

These results demonstrate that SSc-induced remodeling of the fibroblast compartment is sex-specific, with this divergence selectively manifested in profibrotic and inflammatory fibroblast states, most notably through the male-biased expansion of the contractile *COL11A1⁺* lineage.

### Trajectory analysis reveals sex-biased fibroblast-to-myofibroblast transitions

To dissect fibroblast-state transitions across healthy, non-lesional, and lesional skin, we reconstructed a unified transcriptional trajectory using Monocle3. Consistent with their previously established identity, resident myofibroblast progenitor-like cells expressed *CD34, PI16, PIEZO2,* and *SCARA5*^11,21–23^. We observed co-expression of *PI16, CD34*, and *PIEZO2* in both *SFRP2⁺DPP4⁺* and *COCH⁺* clusters, whereas *SCARA5* was predominantly localized to the *SFRP2⁺DPP4⁺* subpopulation (Fig 5a-b), defining two transcriptionally distinct root states.

**Fig. 5.**
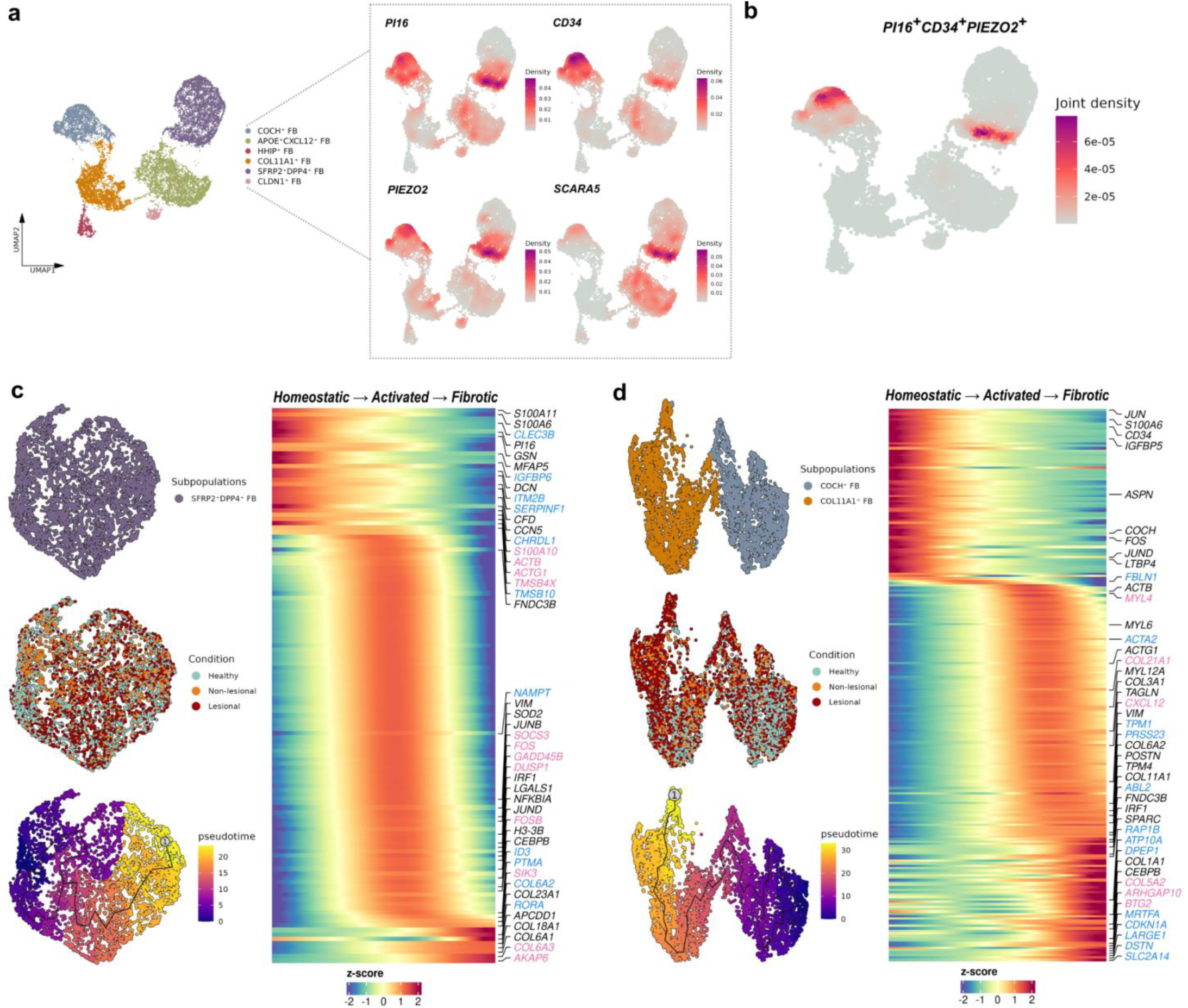
Trajectory analysis and progenitor-to-fibrotic transitions of dermal fibroblasts. **a** UMAP visualization of six fibroblast subtypes (left) with corresponding density plots (right) showing the expression of reported progenitor-associated markers *PI16, CD34, PIEZO2*, and *SCARA5*. **b** UMAP embedding illustrating the joint density of the *PI16⁺CD34⁺PIEZO2⁺* positive cells, identifying a potential progenitor pool. **c-d** Pseudotime trajectory analysis for the *SFRP2⁺DPP4⁺* fibroblast lineage (**c**) and the *COCH⁺–COL11A1⁺* fibroblast lineage (**d**). Left panels show UMAPs coloured by subpopulation identity, skin types, and calculated pseudotime. Right heatmaps display the expression of top dynamic genes that significantly change along the homeostatic–activated–fibrotic transition (Moran’s *I* > 0.3, *q* < 0.05). Colour intensity represents the *Z*-score of gene expression. Annotated genes are categorized by their sex-specific expression patterns: female-biased (pink), male-biased (blue), or shared across sexes (black).

Pseudotime reconstruction from the *SFRP2⁺DPP4⁺* progenitor pool revealed a terminal differentiation trajectory toward a fibrotic state, characterized by coordinated downregulation of progenitor identity and robust induction of ECM gene expression (Fig. 5c, Supplementary Data 7). Sex-stratified pseudotime-dependent gene expression analysis identified divergent transcriptional programs: female-biased trajectories were enriched for acute stress response and cytoskeletal remodeling, whereas male trajectories prioritized translational machinery and metabolic adaptation.

A parallel trajectory originating from the *COCH⁺* cluster culminated in *COL11A1⁺* myofibroblasts (Fig. 5d, Supplementary Data 8). Along this trajectory, myofibroblast-associated genes, ECM components, and cytoskeletal programs progressively increased in both sexes. In females, matrix-associated genes showed dynamic regulation. Conversely, the male trajectory was characterized by the induction of mechanotransduction signaling, senescence markers, and sustained ATP metabolism. This suggests a more contractile and energetically sustained fibrotic state in males. These data demonstrate that fibrotic progression does not follow a uniform differentiation program but instead arises through sex-dependent rewiring of fibroblast lineage trajectories.

### Sex-biased fibroblast–immune cell communication in lesional SSc skin

To elucidate sex-differential interactions between fibroblasts and immune cells in lesional skin, we applied CellChat to infer ligand-receptor-mediated communication networks. Global network analysis revealed sex differences, with females exhibiting higher total communication strength than males (Fig. 6a). While *COL11A1⁺* and *SFRP2⁺DPP4⁺* subpopulations served as major communication hubs in lesional skin in both sexes (Fig. 6b), the magnitude of specific fibroblast–immune cell interactions revealed distinct sex-biased patterns (Fig. 6c). In females, signaling was most prominent between fibroblasts and both conventional type 1 dendritic cells (cDC1) and plasmacytoid dendritic cells (pDCs). Conversely, males showed a signaling bias toward conventional type 2 dendritic cells (cDC2), which exhibited widespread interactions across multiple fibroblast subtypes.

**Fig. 6.**
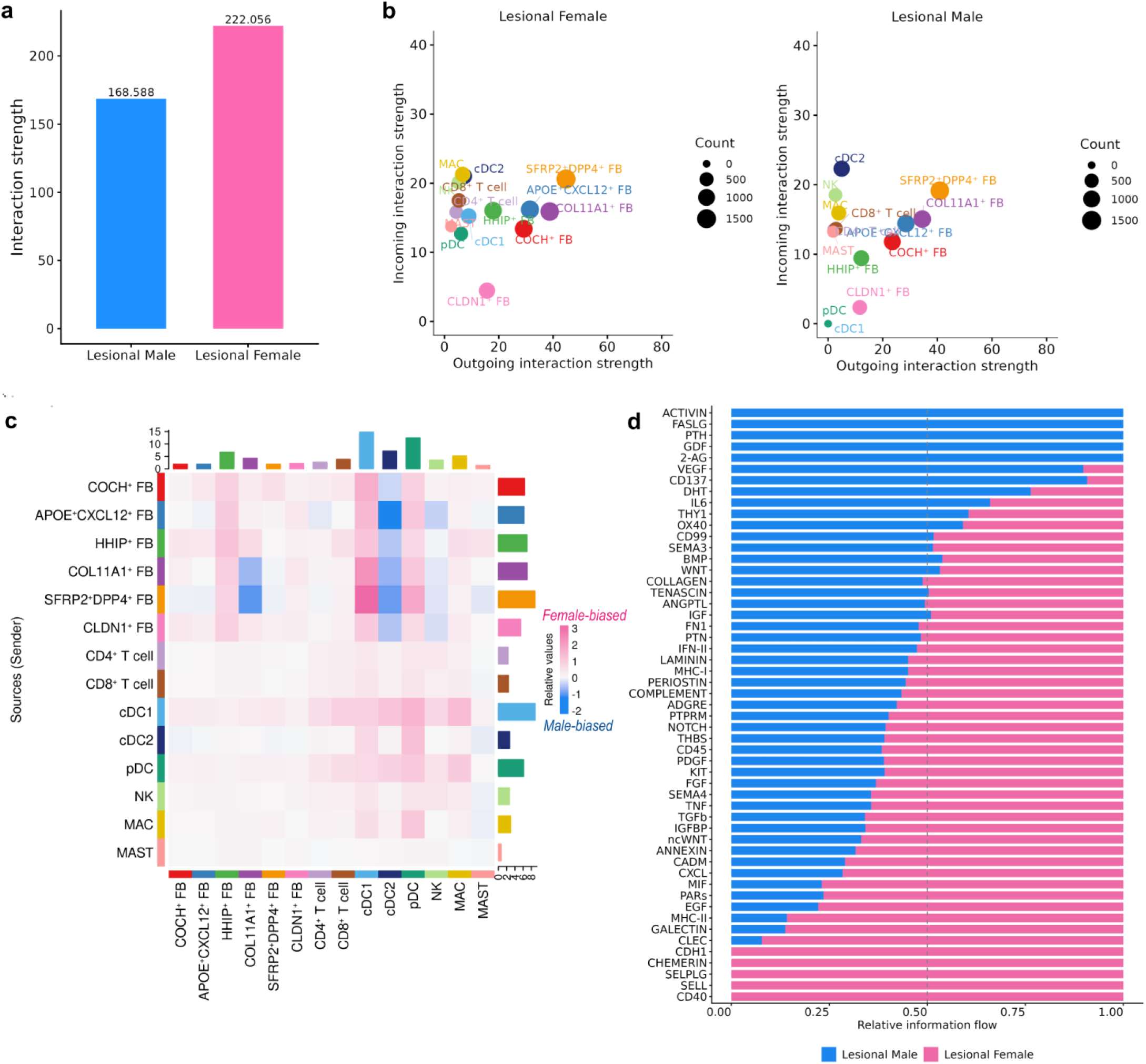
Sex-biased fibroblast–immune cell communication in lesional SSc skin. **a** Bar plot comparison of total cell–cell interaction strength in lesional skin from females (pink) and males (blue). **b** Signaling role scatter plots identifying dominant sender (outgoing) and receiver (incoming) cell populations in lesional skin from females (left) and males (right). Dot size corresponds to the number of inferred interactions. **c** Heatmap showing differential cell–cell communication strength between sender and receiver cell types. Colour intensity represents relative interaction strength, with pink indicating female-biased and blue indicating male-biased interactions. **d** Stacked bar plot showing relative information flow for significant signaling pathways in lesional skin. Pathways are ranked by their relative contribution in females (pink) versus males (blue).

Information flow analysis further identified the signaling pathways driving these sex-biased communication patterns (Fig. 6d). Female lesional fibroblasts preferentially engaged pathways associated with leukocyte recruitment, immune-stromal adhesion, and antigen presentation, including CD40, P-selectin (SELP), L-selectin (SELL), chemerin, major histocompatibility complex class II (MHC-II), and C-type lectin (CLEC) signaling. In addition, the female microenvironment showed enrichment of chromatin regulatory signaling involving chromodomain helicase DNA-binding protein 1 (CHD1), alongside canonical fibrotic and ECM remodeling pathways, such as PDGF, TGF-β, periostin, collagen, and laminin. Male lesional skin was characterized by signaling dominated by growth factor–related pathways, including activin and growth differentiation factor (GDF) signaling, as well as immune co-stimulatory signaling mediated by CD137. Notably, male-biased signaling was characterized by Fas ligand (FASLG)-apoptotic pathways and a shift toward hormonal signaling through the parathyroid hormone (PTH) and dihydrotestosterone (DHT) axes.

## Discussion

Biological sex is a key determinant of SSc pathogenesis, influencing immune activation, fibrosis, and tissue remodeling^3,4,7^, yet the cellular and molecular mechanisms underlying this sex disparity remain poorly understood. Our single-cell transcriptomic analysis of SSc and healthy skin revealed pronounced sex-biased programs in lesion development, fibroblast activation and differentiation, and intercellular communication, contributing to divergent fibrotic outcomes observed between sexes.

The sex-transcriptional asymmetry identified through our interaction model reflects a divergence in the pathological programs governing SSc fibrogenesis. Female SSc fibroblasts showed enhanced activation of immune-inflammatory, collagen-related, and estrogen-responsive pathways. In particular, interferon signaling was significantly enriched in females, consistent with previous findings where it exhibits sex-biased activity^4,5^. Females display broader innate and adaptive immune activity, likely driven by hormonal factors, X-chromosome effects^5,24^, and recently described autosomal sex-differentiated genetic effects^7^, which together may contribute to a more inflammatory skin microenvironment. In this context, estrogens are generally associated with enhanced immune activity, whereas androgens have immunosuppressive effects^4^. In SSc, circulating androgen levels correlate with anti–topoisomerase I autoantibodies, which are more prevalent in diffuse cutaneous subtype^25^, and we observed elevated DHT signaling in male lesional fibroblasts, potentially contributing to increased fibrotic severity. Our findings evidenced stronger inflammatory programs in females and greater fibrotic contributions in males.

Conversely, estrogen has been shown to induce *PDGF* gene expression^26^, providing a potential mechanistic link between hormonal regulation and fibroblast activation. Consistent with this, our results reveal a pronounced female bias in PDGF signaling. Notably, *PDGFB* was also identified as a female-specific susceptibility locus in a recent sex-stratified genomic study of SSc^7^, further supporting a convergence of hormonal and genetic factors underlying sex-dependent differences in fibroblast activation. Concurrent enrichment of PDGF signaling and its downstream KRAS signaling, together with PI3K–AKT–mTOR signaling in female lesional fibroblasts, indicates a heightened capacity for proliferation and collagen deposition. This is consistent with the selective enrichment of the KRAS pathway in fibroblast clusters associated with severe SSc skin and its role in driving fibrosis^27^.

Sex differences in SSc fibrogenesis were associated with divergent activation of TGF-β signaling and downstream cellular programs. Female fibroblasts preferentially engaged canonical TGF-β signaling linked to collagen biosynthesis^28^, whereas male fibroblasts exhibited enrichment of non-canonical TGF-β pathways, including Rho/ROCK, MAPK, and mTORC1 signaling^8,29^, consistent with a more contractile myofibroblast phenotype. This male-biased program was accompanied by increased oxidative phosphorylation and aerobic metabolism, suggesting elevated energetic demands^30^, and may be exacerbated by the absence of estrogen-mediated constraints on excessive myofibroblast differentiation^31^. Concordantly, male fibroblasts showed enrichment of SLIT/ROBO signaling, a pathway linked to mechanotransduction^32,33^. Male fibrogenesis was further characterized by MYC-associated transcriptional programs, previously reported to be overexpressed in SSc skin and implicated in fibroblast activation and fascia-like contractility^34^. In parallel, male fibroblasts displayed enrichment of DNA repair, oxidative stress, and p53-dependent stress response pathways, consistent with accelerated aging and heightened susceptibility to p53-mediated apoptosis^35^. Notably, MYC has also been implicated in modulation of innate immune responses, potentially contributing to reduced interferon signaling and lower inflammatory activity observed in males^36^.

We identified several female-biased TFs in fibroblasts that emerge as key drivers of these responses and likely cooperate with VGLL3, a cofactor promoting female-biased skin autoimmunity and interferon signaling^18^. Furthermore, elevated Hippo pathway activity in female lesional fibroblasts was accompanied by *VGLL3* upregulation, suggesting that VGLL3 may modulate TEAD-dependent transcription downstream of Hippo signaling to sustain pathological matrix remodeling and myofibroblast differentiation in female SSc skin^12^.

Our findings demonstrate that sex-biased programs were compartmentalized within distinct fibroblast subpopulations, primarily *SFRP2⁺DPP4⁺* and *COL11A1⁺* cells. The female-enriched *SFRP2⁺DPP4⁺* population represents a matrix-producing, proto-myofibroblast state characterized by robust collagen synthesis downstream of canonical TGF-β signaling while remaining *ACTA2*-negative^8^, consistent with previously described *COMP⁺* fibrogenic lineages in SSc^37^. In contrast, *COL11A1⁺* fibroblasts, which are preferentially expanded in male lesional skin, exhibit a more mature *ACTA2*-positive myofibroblast phenotype enriched for mechanosensitive programs, resembling contractile mesenchymal states described in fibrotic and keloid tissue^19^. Previous studies reported that the abundances of profibrotic *COMP⁺* and *COL11A1⁺* fibroblasts positively correlate with MRSS and are higher in dcSSc than in lcSSc^37^.

Trajectory analysis further supports sex-dependent rewiring of fibroblast differentiation during fibrotic progression, revealing two distinct routes to the fibrotic state originating from *SFRP2⁺DPP4⁺* and *COCH⁺* progenitor-like cells. Owing to the inherent heterogeneity and plasticity of the fibroblast lineage, these resident cells are dynamically activated under fibrotic conditions to differentiate toward myofibroblasts or alternative mesenchymal fates^38^. The origin cells in *SFRP2⁺DPP4⁺* cluster overlaps with previously described *SFRP2^hi^WIF1⁺* and *SFRP2⁺PI16⁺*progenitors, the latter known to be highly responsive to matrix stiffness^11,21^. Although both trajectories are present in both sexes, the *COCH⁺-COL11A1⁺* axis is preferentially enriched in male lesional skin. *COCH⁺*fibroblasts have been reported to be reduced in SSc skin^37^. Consistent with this, our trajectory analysis suggests that this apparent reduction may reflect progression along a differentiation continuum toward *COL11A1⁺* fibroblasts rather than depletion of this population. This transition may be linked to mechanosensitive activation associated with cochlin and tissue stiffness^30,39^. In support of a pathogenic role, *COL11A1⁺* fibroblasts are enriched in lesional skin and positively correlate with MRSS^37^. The enrichment of this axis specifically in male lesions suggests a self-sustaining feed-forward loop, where tissue stiffening drives the continuous activation and contraction of myofibroblasts^30^, potentially accounting for the more aggressive fibrotic remodeling observed in male patients.

At the level of stromal-immune crosstalk, female lesional fibroblasts preferentially interacted with cDC1s and plasmacytoid DCs—associated with heightened interferon activity in female SSc^4,5^—whereas male fibroblasts showed enhanced communication with cDC2s, a subset implicated in pro-fibrotic remodeling in SSc^40^. The identification of *BCL11A* as a male-specific SSc risk locus^7^, together with its role in regulating dendritic cell differentiation^41^, supports a genetic contribution to sex-biased immune-stromal interactions that may underlie divergent fibrotic trajectories in SSc.

In summary, we identify a fundamental sex-transcriptional asymmetry in SSc skin that drives distinct fibroblast states. Female-biased programs are characterized by canonical TGF-β signaling, ECM biosynthesis, and inflammatory signaling. In contrast, male-biased programs preferentially engage non-canonical TGF-β pathways, mechanotransduction, contractility, and MYC-associated metabolic stress, supporting a more severe fibrotic phenotype. Our findings may help improve current clinical management, providing new opportunities to tailor interventions to sex-differential fibrotic mechanisms.

## Methods

### Skin samples

Skin biopsies were obtained from 17 patients with SSc and 8 healthy controls, yielding a total of 42 biopsies. After cell separation and initial QC sample removal for low cell count, we included 26 SSc samples from lesional and non-lesional skin (females: 8 lesional, 8 non-lesional; males: 6 lesional, 4 non-lesional) and 8 non-lesional control biopsies (4 female, 4 male). Skin samples were collected at Hospital Clínico San Cecilio (Granada, Spain) and Hospital Vall d’Hebron (Barcelona, Spain). Written informed consent was obtained from all participants with approval from the Ethic Committee of Research of the Granada Province (CEIM/CEI). The study was conducted in accordance with the Declaration of Helsinki. SSc diagnosis was established according to the 2013 ACR/EULAR classification criteria. Throughout this study, “sex” refers to biological sex, as determined by chromosomal composition. Participant sex was recorded from hospital records, and further verified at the single-cell level using the CellXY (v0.99.0). Donor characteristics are provided in Supplementary Data 9.

### Single-cell RNA sequencing

Skin biopsies were dissociated into single-cell suspensions following previously published protocols^42^. Single-cell RNA-seq libraries were prepared using the 10x Genomics Chromium Single Cell 3’ Gene Expression platform. Raw sequencing reads were aligned to the human reference genome (GRCh38) using Cell Ranger (10x Genomics), and gene expression matrices were generated for downstream analysis.

### Data preprocessing and integration

Seurat (v5.3.0) was used for preprocessing and downstream analysis of scRNA-seq data^43^. Cells with fewer than 200 or more than 6,000 detected genes were excluded. Genes expressed in fewer than 100 cells were also removed. Cells exhibiting a high proportion of mitochondrial (>10%), hemoglobin (>10%), or ribosomal (>40%) gene expression were filtered out, as such profiles typically indicate stressed, dying, or contaminated cells. Ambient RNA contamination was corrected using DecontX^44^, and potential doublets were identified with both DoubletFinder^45^ and scVI-Solo^46^. Only consensus singlets detected by both methods were retained. After quality control, the integrated atlas comprised 81,232 cells, with a median of 2,687 detected genes and 7,113 transcripts per cell.

Counts were normalized using a shifted logarithm transformation with a scaling factor of 10,000. Data integration across samples was performed using Harmony^47^. Principal component analysis (PCA) was conducted on 2,000 highly variable genes. A k-nearest neighbors graph (k = 20) was constructed in PCA space using the first 20 principal components (PCs), and clusters were identified using the Leiden algorithm (resolution = 0.8). Clusters with similar transcriptional profiles were subsequently merged. To ensure that identified clusters represented reproducible biological states rather than patient-specific effects, we only retained clusters containing cells from at least five independent samples. Two-dimensional visualization was performed using Uniform Manifold Approximation and Projection (UMAP).

Cell-type annotation combined automated classification and manual curation. CellTypist (v1.7.1) with the pre-trained human_skin model was applied for automated cell identity prediction^48^. Cluster markers were identified using Seurat’s FindAllMarkers function (Wilcoxon rank-sum test) and compared with established marker genes from published human skin single-cell datasets^14,15^. Fibroblasts (n = 14,832), defined by expression of *DCN, COL1A1, COL1A2, LUM,* and *PDGFRA,* were subset for downstream analyses.

### Differential expression analysis

DEGs were identified using a pseudobulk approach with edgeR^49^. For each donor, raw counts were aggregated across fibroblasts to generate pseudobulk profiles, which were normalized using the trimmed mean of M-values with singleton pairing (TMMwsp). To capture the full spectrum of sex bias in fibrotic regulation, we performed two complementary analyses. First, we used a generalized linear model with an interaction term between sex (female and male) and skin type (lesional, non-lesional, and healthy) to test whether the magnitude of transcriptional changes associated with lesion development differed between sexes. Immunosuppression status was included as a covariate. To account for unwanted sources of variation, we applied RUVr from the RUVSeq package^50^ to estimate factors of unwanted variation from model residuals (*k* = 15), which were included in the model as additional covariates. Male and healthy skin were set as reference levels. We tested the interaction contrast: *Sex_Female_:SkinType_Lesional_ − Sex_Female_:SkinType_Non-lesional_.* This contrast captures sex differences in the transcriptional transition from non-lesional to lesional skin. Statistical significance was assessed using edgeR’s quasi-likelihood F-test. The logFC reflects the estimated effect of this contrast: positive values indicated a stronger transcriptional shift from non-lesional to lesional skin in females, whereas negative values indicated a stronger shift in males.

Second, we conducted sex-stratified differential expression analyses comparing lesional and non-lesional skin to determine the directionality of fibrotic response. Here, logFC represents the lesional versus non-lesional change within each sex. Genes with FDR < 0.05 and |logFC| > 0.50 were considered significant in all the analyses. Including lesional, non-lesional, and healthy skin in the model, enabled robust estimation of baseline expression and partially accounted for intra-individual variation, with healthy skin serving as the reference.

### Pathway enrichment analysis

To capture sex-biased transcriptional programs in fibrogenesis, we performed GSEA on gene lists derived from differential expression analyses and ranked by the product of sign(logFC) × −log₁₀(p-value). Analysis was conducted using clusterProfiler (v4.18.2)^51^ with the human Molecular Signatures Database, including Hallmark (H) and curated Reactome pathways (C2) collections. Normalized enrichment scores (NES) were used to quantify pathway activity. In the interaction analysis, positive NES indicated female-biased pathways during lesional development, while negative NES represented male-biased programs. In sex-stratified analyses, positive NES indicated pathways upregulated in lesional skin, and negative NES indicated pathways downregulated in lesional skin.

To identify pathway enrichment across fibroblast subtypes, over-representation analysis (ORA) was performed on cluster-specific marker genes (adjusted p-value < 0.05; logFC > 0.5), which were identified via Seurat FindMarkers. Subpopulation-specific pathway enrichment was assessed using the compareCluster function with Reactome pathway annotation. For both GSEA and ORA, pathways with p-value < 0.05 were considered significant.

### Transcription factor activity and pathway activity inference

TF and pathway activities in fibroblast subtypes were inferred from SCTransform-normalized gene expression using the decoupleR framework. TF–target interactions were defined using the CollecTRI network^52^, and TF enrichment scores were calculated using a univariate linear model. To infer sex-biased regulatory signatures, TF activities were inferred from gene lists ranked by sign(logFC) × −log₁₀(p-value) from interaction-based differential expression analysis. Positive TF activity scores indicated female-biased TF activity, whereas negative scores indicated male-biased activity. Additionally, potential interactions between sex-biased TFs and VGLL3 were predicted using TFLink^53^. Pathway activities of fibroblast subpopulations were quantified via PROGENy scores^54^ computed from aggregated SCTransform-normalized expression profiles using a multivariate linear model.

### Fibroblast program scoring

Module scores for fibroblast functional programs including ECM, contractile, and myofibroblast signatures, were calculated across fibroblast subpopulations, sexes, and skin types using Seurat’s AddModuleScore function. Gene sets were derived from Gene Ontology (GO) annotations obtained via msigdbr (C5 collection) and further refined to retain biologically relevant terms. The myofibroblast score was generated by integrating GO gene sets associated with myofibroblast differentiation and key profibrotic pathways, including TGF-β, PDGF, Rho/SMAD, and Hippo signaling pathways^8^, together with previously reported canonical myofibroblast markers^14^. Module scores were z-scaled across cells.

### Differential abundance analysis

Differential abundance of fibroblast subtypes across sexes and skin types was assessed using MiloR (v2.4.1)^55^. Samples with fewer than 40 fibroblasts were excluded to maintain the robustness of neighborhood abundance estimations. A k-nearest neighbor graph (k=30) was constructed based on the top 20 PCs, with cells aggregated into representative neighborhoods (proportion = 0.2). Neighborhood-level cell counts were modeled using a negative binomial mean–variance framework implemented in MiloR, followed by quasi-likelihood F-tests with TMM normalization. To assess sex-biased differential abundance, contrasts were defined within each condition (healthy and lesional skin) comparing females and males, where positive logFC indicated enrichment in females and negative values indicated enrichment in males. Statistical significance was determined using a spatial FDR threshold of 10%. Differentially abundant neighborhoods were annotated by their constituent fibroblast subtypes and visualized using beeswarm plots and neighborhood graph representations.

### Pseudotime trajectory analysis

Trajectory inference was performed using Monocle3^56^, with the *COCH^+^*-*COL11A1^+^*and *SFRP2^+^DPP4^+^* compartments analyzed separately. Trajectories were rooted in a progenitor-like state characterized by the expression of *PI16, CD34, PIEZO2*, and/or *SCARA5*. To identify sex-biased genes associated with the fibrotic transition, spatial autocorrelation was assessed using graph_test function on the integrated dataset (males and females combined) and independently within each sex. Genes exhibiting dynamic expression along the trajectory were defined by a Moran’s *I* > 0.3 and q-value < 0.01. Shared dynamic genes were identified as the intersection of significant genes, whereas sex-specific dynamic genes were defined by set differences.

### Cell-cell communication analysis

Fibroblast and immune cell clusters were merged into a single object, and cell-cell communication was analyzed using CellChat (v1.6.1)^57^. To ensure balanced comparisons, female and male lesional fibroblasts and immune cells were downsampled to equal numbers of cells prior to analysis. Lesional male and female datasets were analyzed separately to construct independent signaling networks. Sex-biased communication was assessed using CellChat’s comparative framework by contrasting independently inferred male and female networks. Sex bias in cell communication strength was quantified as differences in inferred interaction strength between sexes. In addition, sex-biased signaling pathways were identified by comparing global information flow, defined as the sum of communication probabilities across all cell-cell pairs within each signaling pathway.

## Acknowledgements

We thank Sofia Vargas and Gema Robledo for their exceptional technical expertise. We also thank all of the participants who provided the data for their essential collaboration. This work was supported by the Instituto de Salud Carlos III (PI22/00092 [co-financed by the EU], Redes de Investigación Cooperativa Orientadas a Resultados en Salud [RD24/0007/0016]), and by grant PID2022-139292OB-I00 funded by MICIU/AEI/10.13039/501100011033 and by ERDF/EU. CK is beneficiary of a Juan de la Cierva 2023 postdoctoral fellowship (JDC2023-052379-I) financed by MICIU/AEI/10.13039/501100011033 and by the FSE+. IR-M was supported by the programme Formación de Profesorado Universitario (FPU21/02746) funded by the Spanish Ministry of Science, Innovation, and Universities (MICIU). This research is part of the doctoral degree awarded to IR-M within the Biomedicine programme from the University of Granada.

## Author contributions

MA-H and JM contributed to the conception and study design. CK, EA-L, MA-H, and MK contributed to data analysis, visualization, and interpretation. JLC, NO-C, AG-D-C, CPS-A, and RR-V contributed to sample collection. IR-M and GV-M contributed to sample processing. CK, MA-H, JM, IR-M contributed to manuscript drafting. All authors participated in manuscript revision and approved the final version of the manuscript.

## Data availability

The single-cell RNA-seq data generated in this study will be deposited in the Gene Expression Omnibus (GEO) database upon publication. All other data are available in the article and its Supplementary files or from the corresponding author upon reasonable request.

## Code availability

All the scripts used in this study for the analysis are deposited in GitHub.

