## Supplementary Fig for "Sex-biased Fibroblast Subpopulations and Transcriptional Programs Reveal Mechanisms of Skin Lesion Development in Systemic Sclerosis"

**Supplementary fig 1.** Sex-stratified differential expression analysis

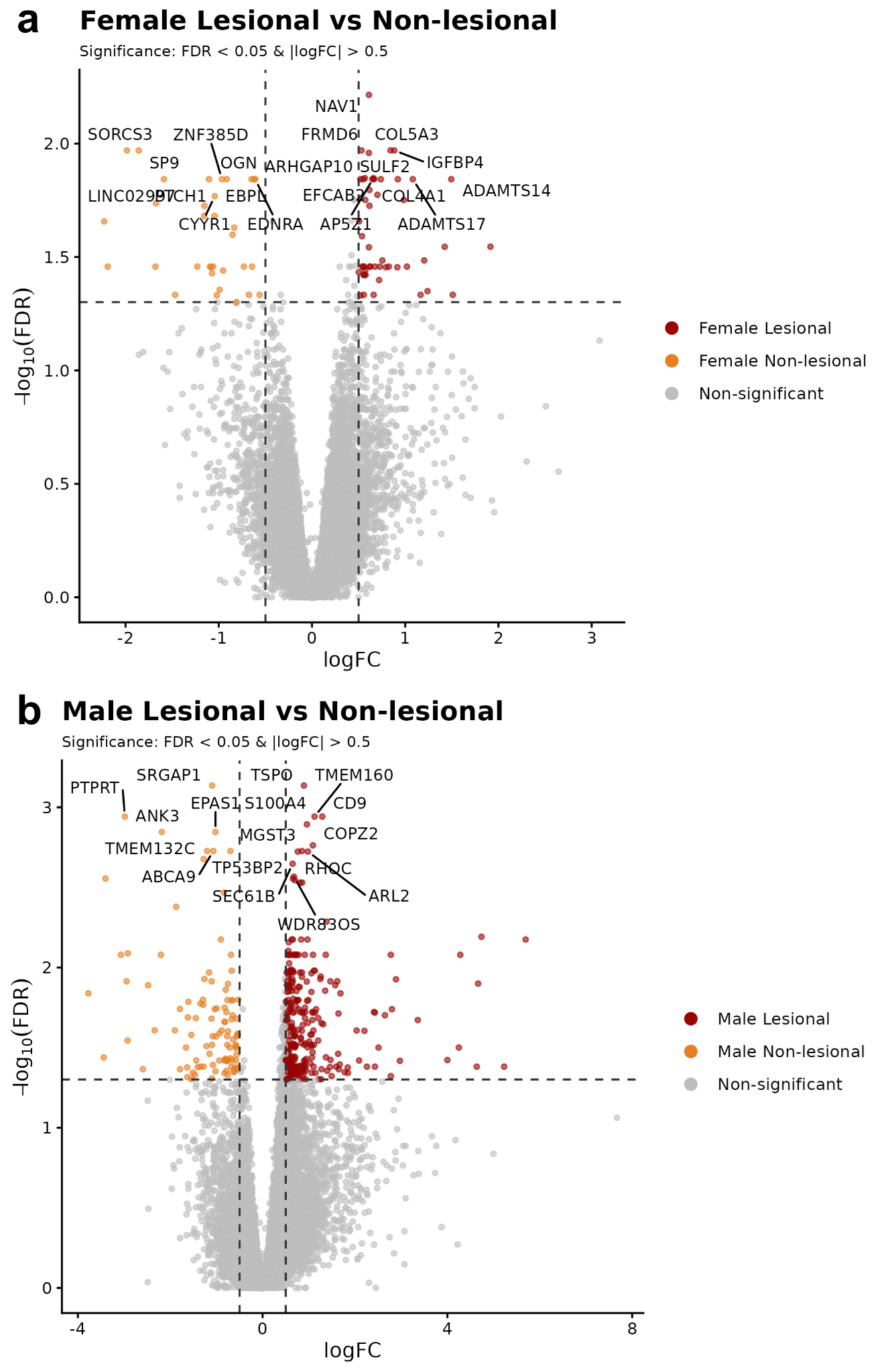

Volcano plots comparing gene expression profiles in (**a**) females and (**b**) males between lesional and non-lesional skin. Genes meeting the significance threshold (FDR < 0.05, |logFC| > 0.5) are highlighted to show lesional (red) and non-lesional (orange) bias.

**Supplementary fig 2.** VGLL3 interaction landscape and expression across fibroblast subpopulations

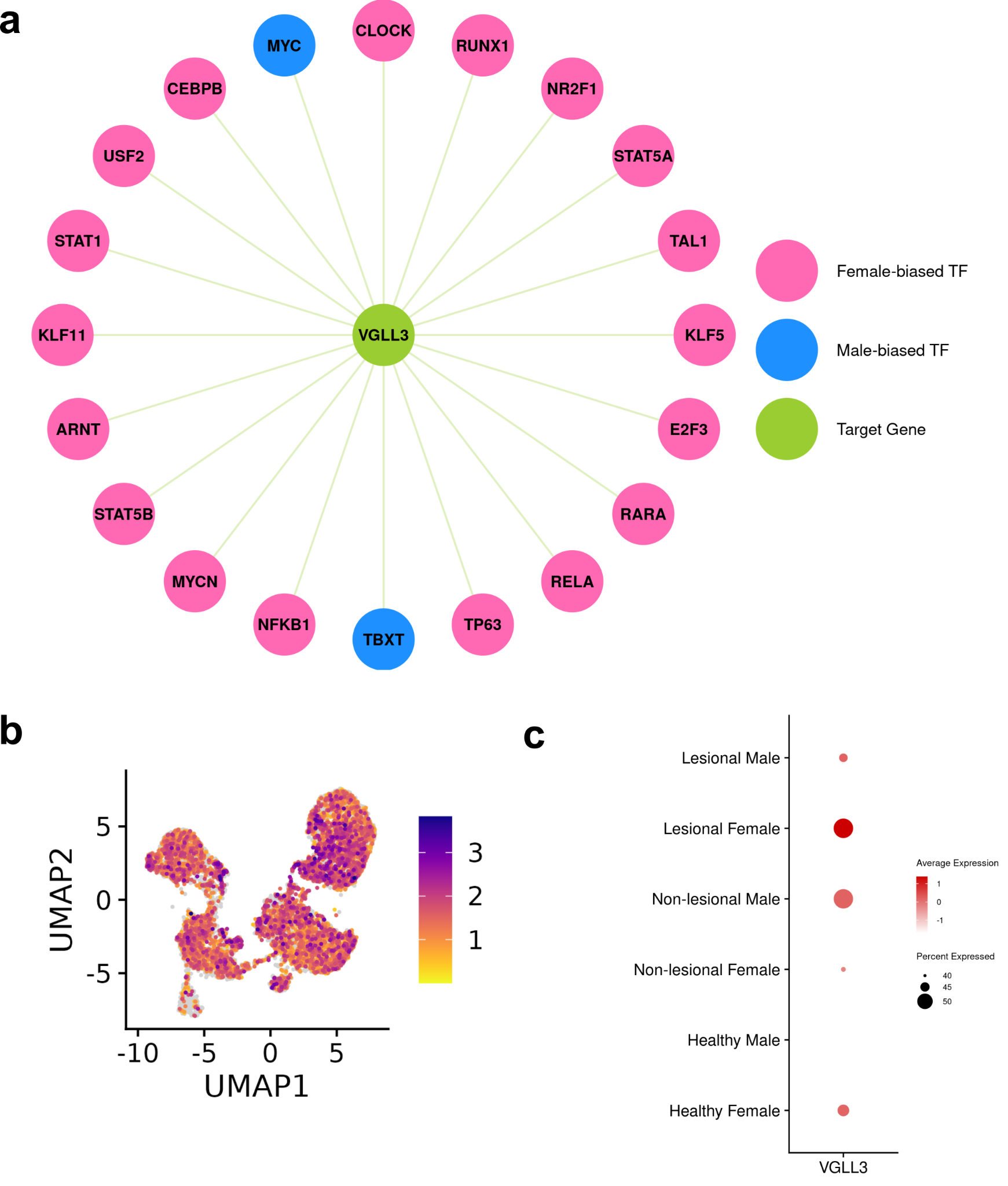

**a** Predicted interactions between sex-biased transcription factors and VGLL3 identified using TFLink. **b** UMAP visualization of *VGLL3* expression in the fibroblast subpopulation. **c** Dot plot showing the average expression and percentage of *VGLL3*-expressing cells across sexes and skin types.
